# Coding translational rates: the hidden genetic code

**DOI:** 10.1101/062935

**Authors:** Luis Diambra

## Abstract

In this paper we propose that translational rate is modulated by pairs of consecutive codons or bicodons. By a statistical analysis of coding sequences, associated with low or with high abundant proteins, we found some bicodons with significant preference usage for either of these sets. These usage preferences cannot be explained by the frequency usage of the single codons. We compute a pause propensity measure of all bicodons in nine organisms, which reveals that in many cases bicodon preference is shared between related organisms. We found that bicodons associated with sequences encoding low abundant proteins are involved in translational attenuation reported in SufI protein in *E. coli.* Furthermore, we observe that the misfolding in the drug-transport protein, encoded by *MDR1* gene, is better explained by a big change in the pause propensity due to the synonymous bicodon variant, rather than by a relatively small change in the codon usage. These findings suggest that bicodon usage can be a more powerful framework to understand translational speed, protein folding efficiency, and to improve protocols to optimize heterologous gene expression.

## Introduction

The central dogma of the molecular biology establishes that the information that specifies which amino acid monomers will be added next during protein synthesis is coded in one or more nucleotide triplets known as codons^1^. The genetic code establishes a set of rules that associate the 20 amino acids and a stop signal with 64 codons. This code is almost universal with few exceptions^2^. As there are more codons than encodable signals (amino acids and stop signal) the genetic code is considered degenerated. However, it is well known that synonymous codons are not used with the same frequency. The biased codon usage is a pervasive feature of the information encoded in genomes, but it is not universal because different species have different associated preferences^1^. The existence of selective pressures to promote the codon usage bias highlights the complex nature of synonymous codons choices^3,4^. Early reports have pointed out that in prokaryotes the bias is towards codons with high translation rates^5,6^. In this sense, Guimares *et al.* established that elongation rate is affected by the specific amino acid composition, as well as by codon bias, in *E. coli*^7^. On the other hand, the impact of codon usage on translational rates in eukaryotes, where the mRNA processing can also affect the overall translational rate, is an active topic of research^8–13^. However, the role of codon usage has gone beyond the translational rates because new experimental findings suggest that codons with slow translation rates temporally separate the synthesis of defined protein portions and can synchronize the synthesis with the concurrently folding process of the proteins domains^14–17^. It has been proven that translational pauses can schedule the sequential folding schemes and can lead to different protein conformations^17^, and that the functionality of translated proteins can be affected by replacing rare codons with more frequently used codons^18–20^. In addition to the use of rare codons associated with scarce tRNA usage, there exist other mechanisms to modulate the speed of translation or to cause pauses. Among them, we can mention the blocking of ribosomal transit due to secondary structure elements in mRNAs^21^, and interactions of basic residues in the nascent polypeptides with the wall of the ribosomal exit tunnel^22^. Furthermore, Li *et al.* showed that translational pauses in *E. coli* are coded by sequences similar to the Shine-Dalgarno sequence^23^.

However, in the last years emerging evidence has shown that the translational rate could be encoded by a sequence longer than a triplet, in particular by pair of consecutive codons, hereafter, bicodons^24^. In this sense, a study on over 16 genomes has revealed that bicodons formed by two rare codons are frequently found in prokaryotes but rarely used in eukaryotes^25^. In addition bicodons such as NNUANN are universally underrepresented, whereas NNGCNN bicodons are mostly preferred^26^. More recently, it was reported that rare arginine codons, followed by proline codons, were among the slowest translated bicodons^27^. This evidence could be consequence of the codon co-occurrence bias mechanism^28,29^or the kinetics of the mRNA translocation from the A-site to the P-site^30^. Codon pair bias was also observed in several viral genomes, which matched the codon usage bias of the host^31^. This fact has been used to produce synthetic viruses with attenuated virulence as a new strategy for vaccine development^32^.

Thus, coding sequences seem to carry further information than the information strictly needed for specifying the linear sequence of amino acids in the protein. This additional information is linked with the overall rate of synthesis of the associated protein and the pauses required for the acquisitions of its correct native structure. Despite the enormous impact that this subliminal coding on biotechnology, there are few systems biology methods to associate nucleotide sequences with the rate of protein synthesis^13^. Among them, we can mention the sequencing of ribosome-protected mRNA fragment or ribosome profiling. This methodology has been used to correlate mRNA levels with codon decoding times^33^. In this paper, we present an alternative manner to identify coding sequences that can modulate the ribosomal transit on the mRNA. In this comprehensive survey we did a statistical analysis of bicodon usage frequencies over two sets of proteins, the low abundant and the high abundant proteins, across nine organisms. Our main finding is that there is an important bias of the bicodon usage depending on the protein abundance. In this sense, we determine which bicodons are statistically associated with low or high translational rates, and in which cases such bias can be explained or not by the codon usage bias. Furthermore, we present suggesting evidence for the role of bicodons in the coding translational rate in two well studied cases. In the first case, we show that there exist clusters of bicodons related to low abundant proteins, associated with ribosomal pauses in the sufIsynthesis in *E. coli*^15^. In addition, we also found that the alteration in the structure and function of the MDR1 protein^16^ associated with a synonymous single polymorphism can be better explained by a relatively big change (around 200%) in pause propensity than by a moderate change in codon usage (around 30%).

## Results

### The preferences of the bicodons

The aim of this paper is to associate coding sequences with their relative translational speeds. We expect that this fact to be reflected in differences in the frequency of both codons and codon pairs occurrence in coding sequences associated with proteins with high and low abundance. To check this hypothesis, we select a set of 500 coding sequences associated with proteins with highest abundance, and another 500 coding sequences associated with proteins with lowest abundance, in nine model organisms from different kingdoms. Before showing the whole analysis across several organisms, we begin with an illustrative example. Fig. 1A shows the histogram of the bicodons (red bars) which codifies for the amino acid pair KK, obtained from 500 sequences of *S. cerevisiae* with the lowest protein abundance (PA).

**Figure 1.**
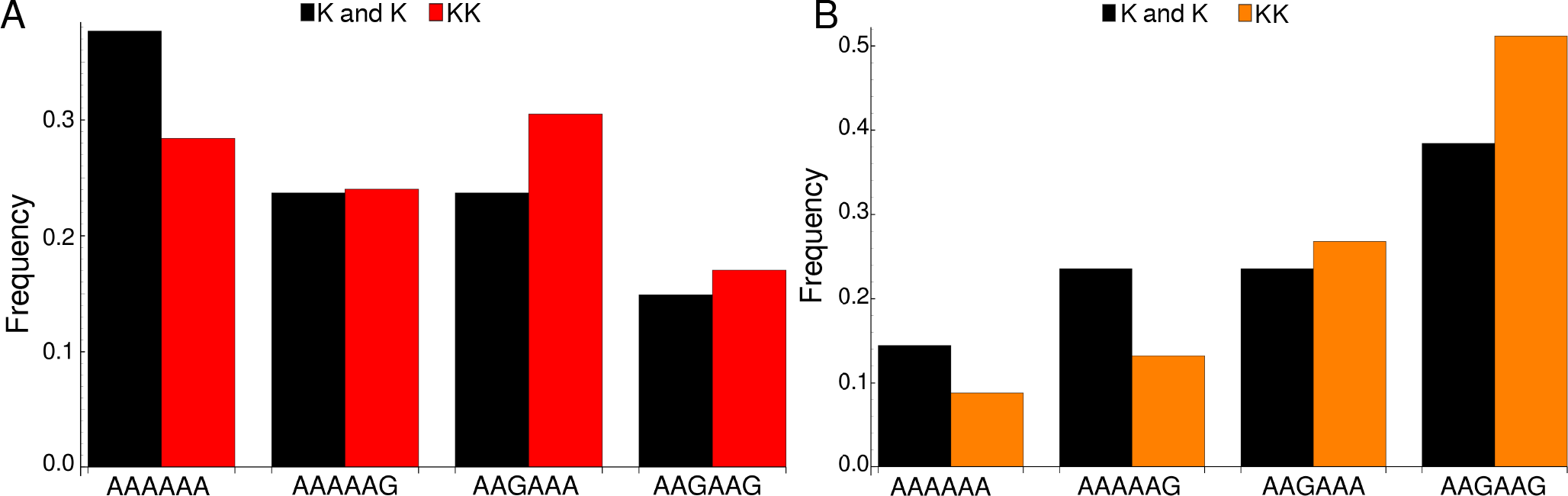
Bicodon usage for the KK amino acid pair. Red and orange bars denote the frequency of bicodons observed in a set of 500 coding sequences with the lowest PA (A), and observed in a set of another 500 coding sequences with the highest PA (B), respectively. Black bars represent the expected frequency obtained by the product of each codon frequency.

We can observe that bicodon usage is not uniform, i.e., it is biased; this fact could be the simple consequence of the known bias observed at the codon level. However, the expected frequency associated with such bicodons (black bars, obtained by the product of each codon frequency) shows that, although some bicodon frequencies can be explained by the bias in the codon usage (for example, the bicodon AAAAAG), some other bicodons have an associated usage frequency that is underrepresented (such as the bicodon AAAAAA), or overrepresented (as the bicodon AAGAAA). This means that two consecutive codons used for coding a given amino acid pair can be correlated. A similar analysis can be performed with sequences associated with the highest PA, as shown in Fig. 1B, and in all other amino acid pairs. Evidence for nonrandom associations between codon pairs, even once codon bias and bias against specific amino acid pairings were subtracted, was previously reported in *E. coli*^34^, and across many other genomes^25^.

However, what is a new remarkable fact in Fig. 1 is the strong difference between the histograms computed for the low and high PA samples. Fisher’s exact test allows one to reject, with high significance level, the null hypothesis that bicodons are equally used in sequences from the low and high PA samples. In the particular case of bicodon AAGAAG the *p*-value is 5.3 × 10^−93^. It is important to point out that the two samples of sequences (500 coding sequences for the proteins with the lowest and highest PA) can introduce an additional bias. In this sense, it is known that protein abundance correlates negatively with coding-sequence length in yeast^35^. To go further in our analysis, we subtracted this bias by constructing two new samples of low and high PA but with similar sequence length distribution, as indicated in the Method section. All subsequent analyses will be made with these unbiased samples (the list of coding-sequence in these samples is given in Supplementary Tables S1 and S2). Fig. 2 shows the histograms corresponding to both low (red bars) and high (orange bars) PA sequences from unbiased samples. It can be seen that bicodon AAAAAA is more frequently used in sequences with low PA than in sequence with high PA, while the frequency usage of bicodon AAGAAG has an inverse relationship. For example, in the last case we have computed the *p*-value from the contingency tables of bicodon AAGAAG which is around 6.5 × 10^−48^, less significant than the one obtained for the biased samples. Figure 2 only illustrates the particular case of KK pair in *S. cerevisiae.* In order to see a broader coverage over amino acid pairs and bicodons we have devised two alternative heat maps: (i) the statistical distance between the frequencies associated with bicodons that encodes a given amino acid pair, (like the histograms depicted in Fig. 2), and (ii) the pause propensity *π*, related to the *p*-value of the Fisher’s exact test, see Method section for its definition.

**Figure 2.**
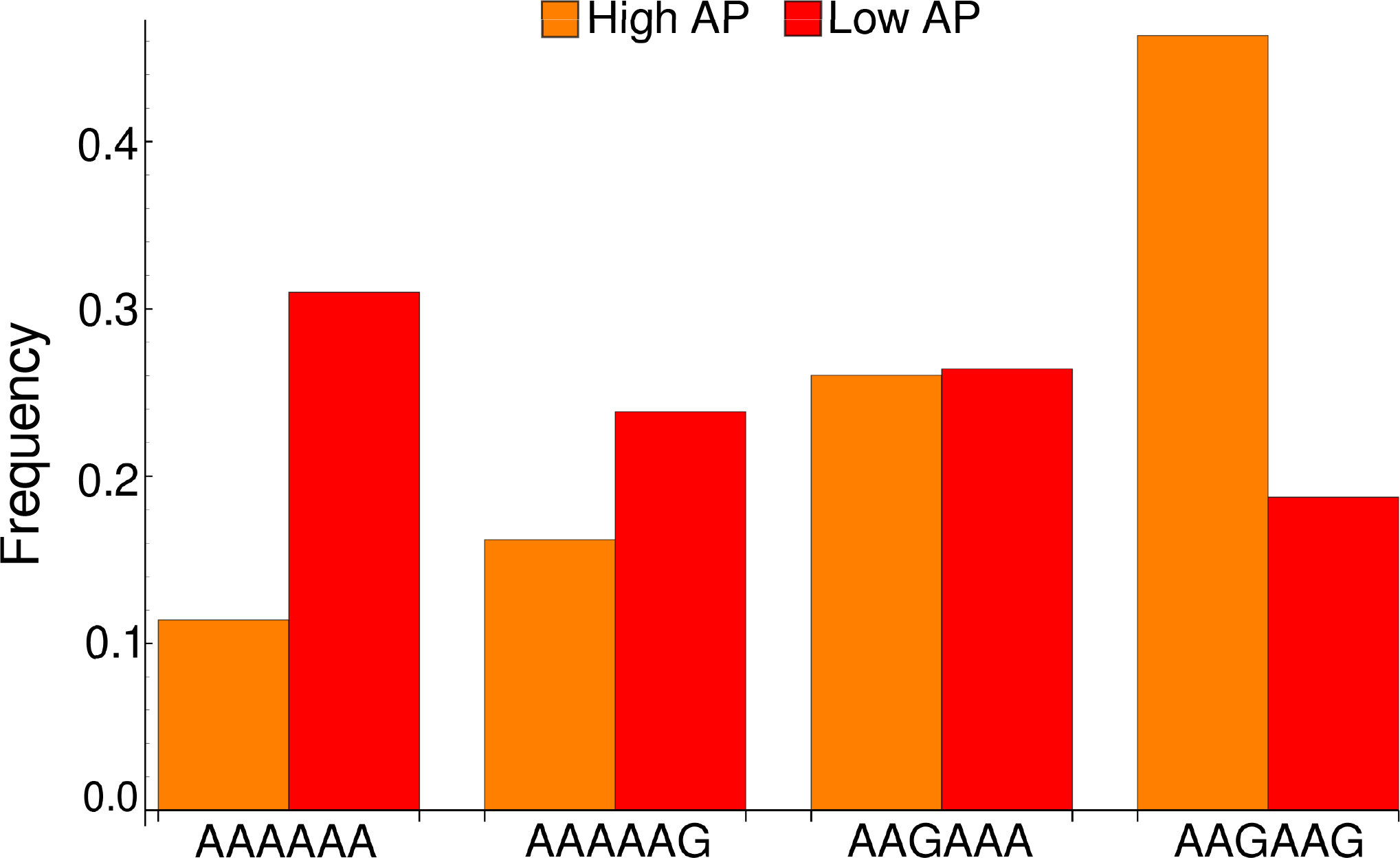
Frequency distributions from unbiased samples of sequences. Frequencies associated with bicodons that encode the amino acid pair KK, computed using sequences from the low PA sample (red bars) and from the high PA sample (orange bars). The frequency usage of bicodon AAGAAA in the sequences of both samples are almost the same, while other bicodons have an evident preference for sequences associated with low or high PA.

### The statistical distance and the pause propensity measures

As statistical distance between the frequency distributions associated with low and high PA samples, we computed the Kullback-Leibler divergence *D_LH_* across all amino acid pairs and all studied organisms. Fig. 3 depicts a heat map (21 P-site codons × 20 A-site codons), where each color pixel represents the quotient *D_LH_/log(n)* for a given amino acid pair, and *n* denotes the number of synonymous bicodons for such dipeptide. A high value (red color) indicates a large discrepancy between the usage frequencies in both samples for a given amino acid pair. This figure shows that the discrepancy in the bicodon usages between low and high PA samples is not the same across organisms, it is a particular feature like the codon bias. It can be seen that *B*. *subtilis* and *S. cerevisiae* have many dipeptides with relatively large divergences, a feature that is also shared by *M. aeruginosa*, *A. thaliana* and both invertebrates, *D. melanogaster* and *C. elegans* (see Supplementary Fig. S1). The studied mammalians (*H. sapiens* and *M. musculus)* and *E. coli* are on the other side. Interestingly, it was in *E. coli,* where bicodon bias was first reported^34^, however, it is clear that bicodon usage bias across the ORFeome does not imply a different usage preference between highly and lowly expressed proteins. Another important feature of the heat maps in Fig. 3 is that they represent a nonsymmetric matrix, i.e., the divergence *D_LH_* of dipeptide X_1_X_2_ is not necessarily equal to the divergence associated with X_2_X_1_. This fact cannot be explained solely with codon usage bias, and reveals a complex correlation between two consecutive codons. Fig. 3 offers a general view of the 420 pairs. However, it can be observed that even when *D_LH_* can give a relatively small divergence, like the one associated with KK pair in *S. cerevisiae,* there could exist one or more bicodons with high difference of occurrence in both samples, such as bicodons AAAAAA and AAGAAG in Fig. 2. To appreciate the differences at a bicodon resolution, the second heat is used. In it, the color of each cell is determined by the pause propensity *π* index, which establishes when the bicodon has preference for sequences with low or with high PA, see Method section for this definition. Fig. 4 shows the heat map associated with all organisms. In order to put in evidence some trends or rules across the studied organisms, the colors of the cells were clustered by similarity using an agglomerative method. The pause propensity *π* of all bicodons and organisms is listed in Supplementary Table S3.

**Figure 3.**
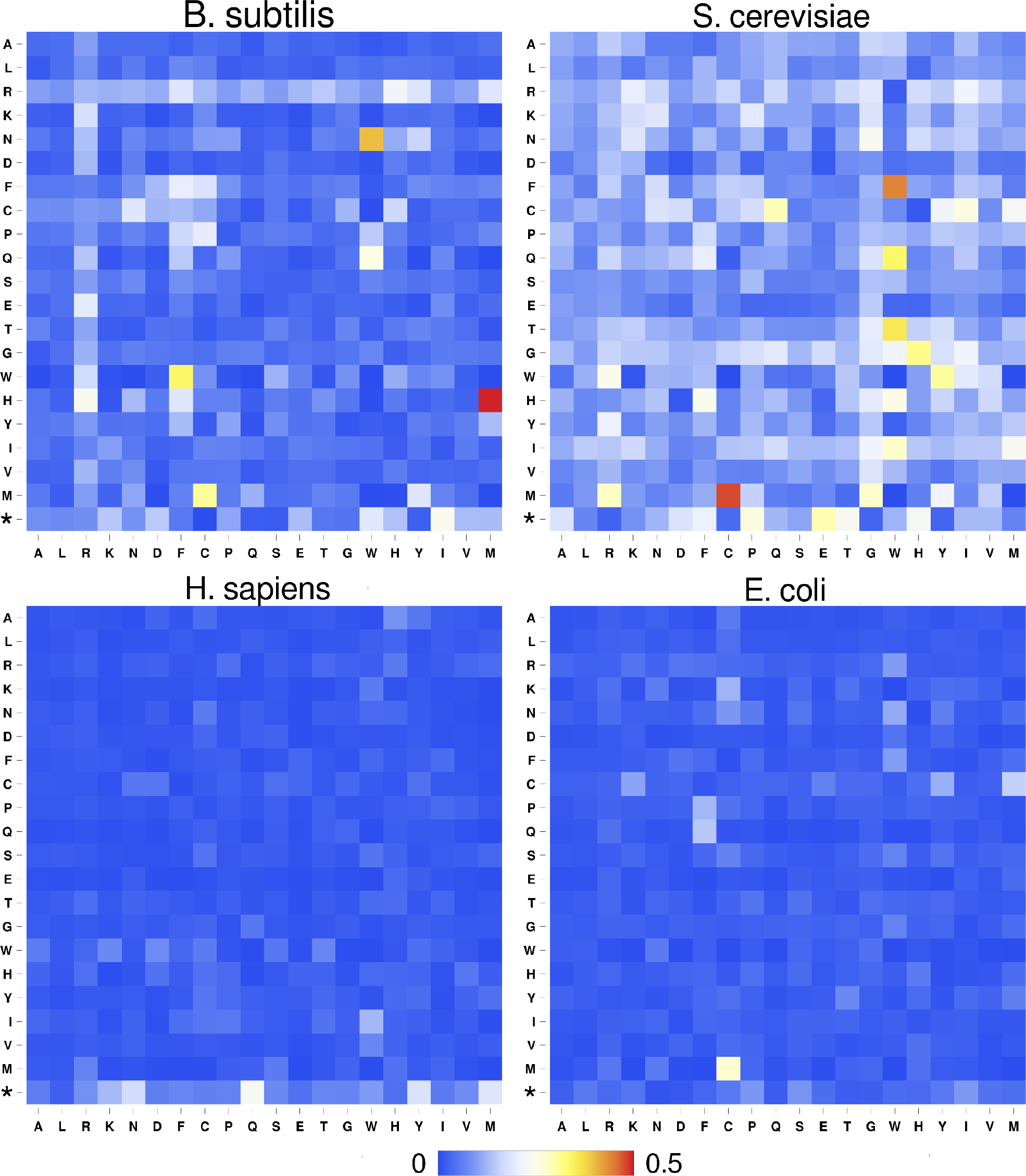
Divergence measure between histograms obtained from low and high PA samples. Statistical distances between all histograms as in Fig. 2 for *B. subtilis*, *S. cerevisiae*, *H. sapiens* and *E. coli.* P-site codons occupy the horizontal axis and A-site codons the vertical axis. Divergence measures associated with other organisms are displayed in Supplementary Fig. S1.

**Figure 4.**
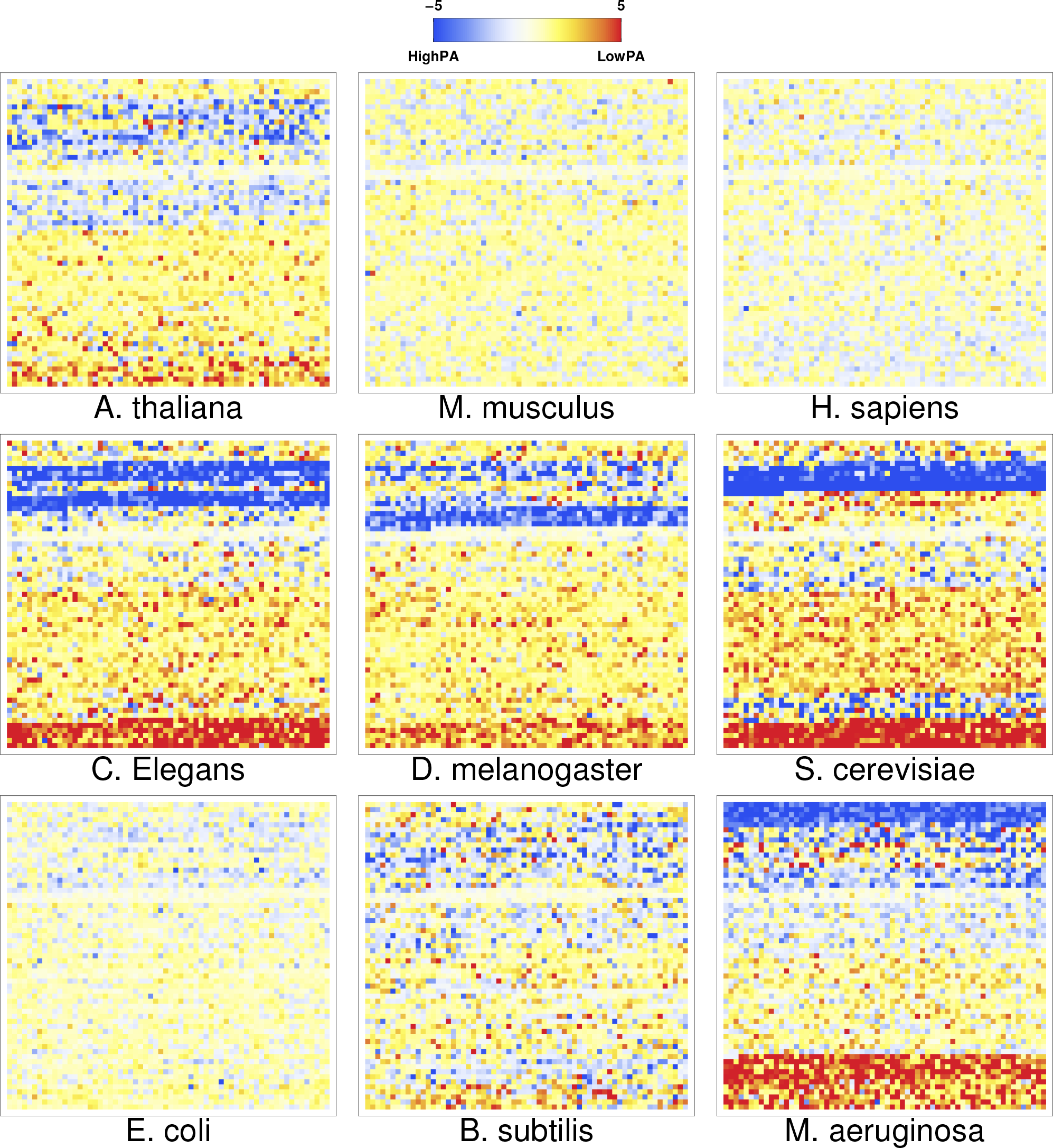
Pause propensity heat maps for nine organisms. The color of each cell is determined by the pause propensity *π* of the associate bicodons. Red cells indicates bicodons with clear preference for sequence associated to low PA, while blue cells indicates bicodons with high PA preference.

From the heat maps of Fig. 4 it can be seen that some bicodons, indicated by blue cells on the top side of the grid, have a clear preference for sequences associated with high PA. In some cases this feature is shared by several organisms such as: *A. thaliana*, *C. elegans*, *D. melanogaster*, *S. cerevisiae*, and *M. aeruginosa.* On the other hand, there are other bicodons, indicated by red cells at the bottom of the grid, which are more frequently used in sequences associated with low PA. Some bicodons have different sequence preferences depending on the organism. For example, there are blue cells on the bottom side of the yeast heat map, that have preference for sequences with low PA in the *M. aeruginosa* heat map. The bicodon preferences are less apparent (less intense colors), and also less frequent, in *H. sapiens*, *M. musculus* and *E. coli.* The low preference of bicodon usage observed in *E. coli* can be a consequence of the fact that there are not a clear distinction between the protein abundance distributions of both samples in this particular case (see Methods). It can be also observed a couple of white rows in all organisms. These cells correspond to bicodons which do not exhibit any preference or they are usually poorly used in both sequence samples and have associated poor statistic.

The heat maps shown in Fig. 4 are very useful to see some common features among organisms, however they do not show whether the bicodon preference is explained or not by the preference of the codon in the pair for sequences associated with low or high PA. In order to study this, we have computed residual scores for each bicodon over sequences with low PA, 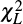, and over sequences high PA, 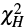. When the residual score is high, the bicodon usage cannot be explained by the codon usage in the same sample of sequences, and has been used previously^25,34^. The value of these residuals, observed frequencies and pause propensity values for all codon pairs and organisms are listed in Supplementary Table S3. In Fig. 5 it can be seen a raster plot of these residual scores for all bicodons in *B. subtilis,* yeast, humans and *E. coli* (residual plots associated with the other five organisms are displayed in Supplementary Fig. S2). As we have two sequence samples for each bicodon it is convenient to take the quantity 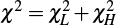 as a whole residual score. To be conservative, we have considered for all organisms that bicodons with *χ*^2^ ≥ 5 are bicodons whose preference is not explained by the preference of the individual codons in the pair.

**Figure 5.**
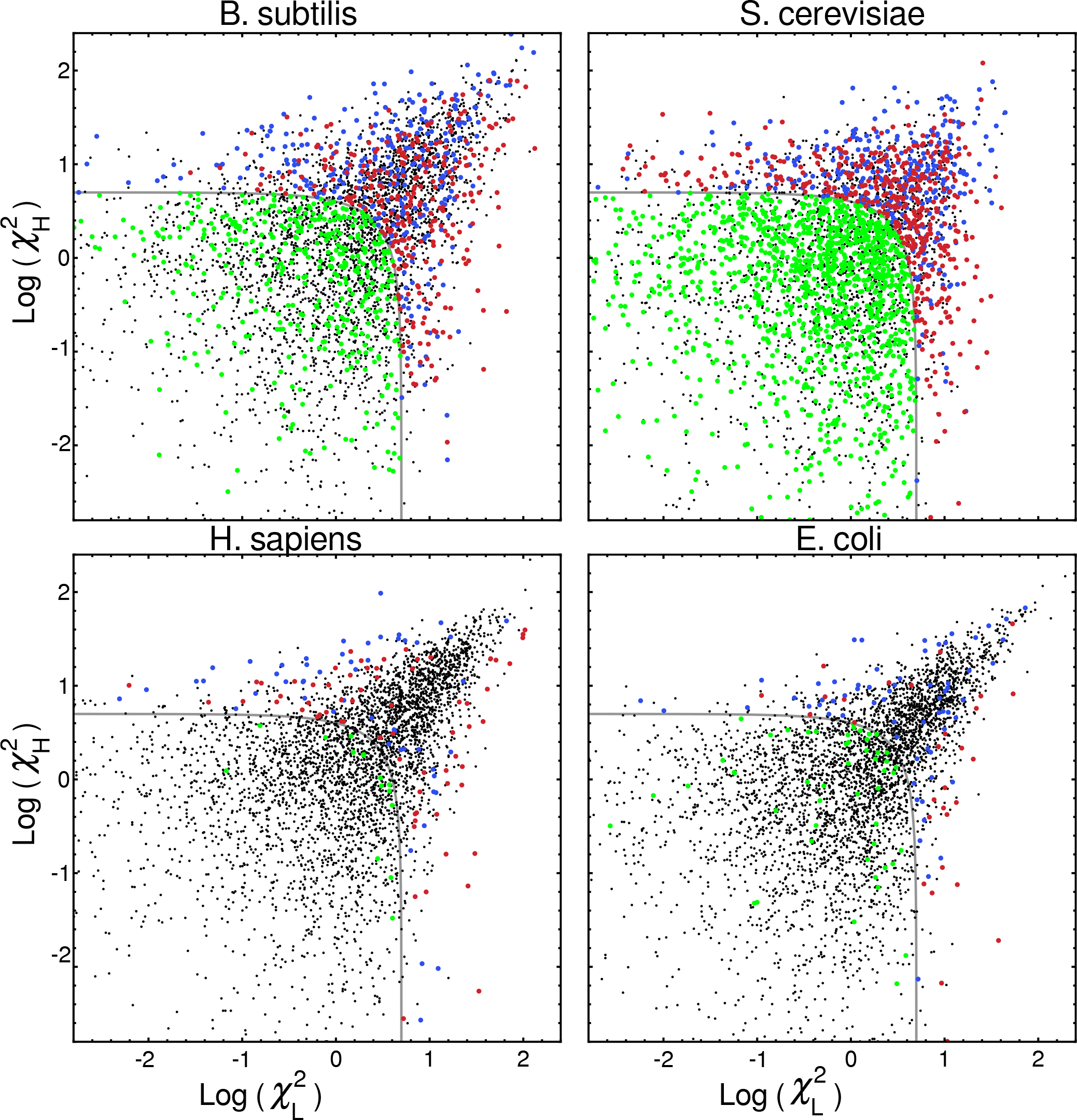
Scatter plots indicating the residual scores *χ_L_* and *χ_H_* computed over the low PA and high PA samples, respectively, for *B. subtilis*, *S. cerevisiae*, *H. sapiens*, and *E. coli*. Raster plots of other organisms are displayed in Supplementary Fig. S2. The codon pairs whose preference for sequences with low or high PA cannot be explained by the codon usage bias are out of the grey quadrant (i.e., *χ*^2^ ≥ 5). Among them, we distinguish bicodons more frequently used in low PA sequences (red dots), or in high PA sequences (blue dots). Inside the quadrant, there are codon pairs with a significantly different usage frequency in low and high PA samples, but whose bias can be explained by codon usage bias (green dots). Codon pairs whose usage frequencies in low and high PA samples are not significantly different (black dots).

Thus, four types of bicodons on the raster plot of Fig. 5 can be distinguished: (i) codon pairs that are significantly more used in sequences associated with low PA than in sequences associated with high PA (*p*-value ≥ 2), and whose preferences cannot be explained by the codon usage bias (red dots); (ii) codon pairs which are significantly more used in sequences associated with high PA than in sequences associated with low PA (*p*-value ≥ 2), and whose preferences cannot be explained by codon usage bias (blue dots); (iii) codon pairs with a significantly different usage frequency in low and high PA samples, but whose preferences can be explained by the codon usage bias, i.e., *χ*^2^ < 5 (green dots), and finally (iv) codon pairs whose usage frequencies in low and high PA samples are not significantly different, i.e., *p*-value ≥ 2 (black dots).

These plots indicate that while there are many bicodons with evident preference for low and high PA sequences in *B. subtilis* and *S. cerevisiae*, there are only few in *H. sapiens* at this significance level. As in humans, mouse and surprisingly also *E. coli* have few bicodons with evident preference (Supplementary Fig. S2). However, in an example below we will see that synonymous SNP associated with changes in the bicodon preference can explain documented protein misfolding and pathological condition in humans. On the other hand, *S. cerevisiae* has more codon pairs than *B. subtilis*, with a significant different usage frequency in low and high PA samples, but such bias is explained by the codon usage bias. The plots for *C. elegans* and *D. melanogaster* (Supplementary Fig. S2) are similar to the raster plot obtained for yeast. In particular these organisms share many bicodons with the same preference listed in Table 2.

**Table 1.** Shared bicodons in *C. elegans*, *D. melanogaster* and *S. cerevisiae* that have high preference for low PA or high PA sequences (*p*-value ≥ 3), but such preference cannot be explained by the codon usage bias (*χ*^2^ > 5).

| dipeptide | bicodon | $\chi_L^2 + \chi_H^2$ | | | $-\log_{10}(p\text{-value})$ | | |
| --- | --- | --- | --- | --- | --- | --- | --- |
|  |  | elegans | fly | yeast | elegans | fly | yeast |
| NA | AATGCA | 8.26623 | 7.0634 | 23.875 | 4.5709 | 24.5975 | 4.0672 |
| NA | AATGCG | 12.6365 | 4.2896 | 13.832 | 3.1346 | 5.55667 | 4.1022 |
| LR | CTTCGA | 44.9244 | 5.3305 | 5.69759 | 4.1911 | 5.24396 | 5.4465 |
| KK | AAAAAG | 47.0125 | 4.9035 | 6.67079 | 8.2823 | 34.0884 | 6.1609 |
| YK | TATAAA | 17.1832 | 4.0035 | 8.81229 | 5.2348 | 5.06749 | 6.0497 |
| FF | TTTTTT | 8.103 | 13.5285 | 18.1809 | 3.7197 | 7.50482 | 12.1921 |
| FQ | TTTCAG | 11.1061 | 7.903 | 7.37717 | 5.2912 | 5.7149 | 6.4614 |
| IT | ATAACA | 11.1095 | 8.6769 | 6.45525 | 3.333 | 5.36998 | 11.9294 |
| KI | AAAATA | 8.20946 | 21.1178 | 56.1534 | 4.691 | 12.958 | 26.2017 |
| IM | ATAATG | 14.6726 | 7.9497 | 6.75161 | 7.0112 | 5.42947 | 4.3485 |

**Table 2.**
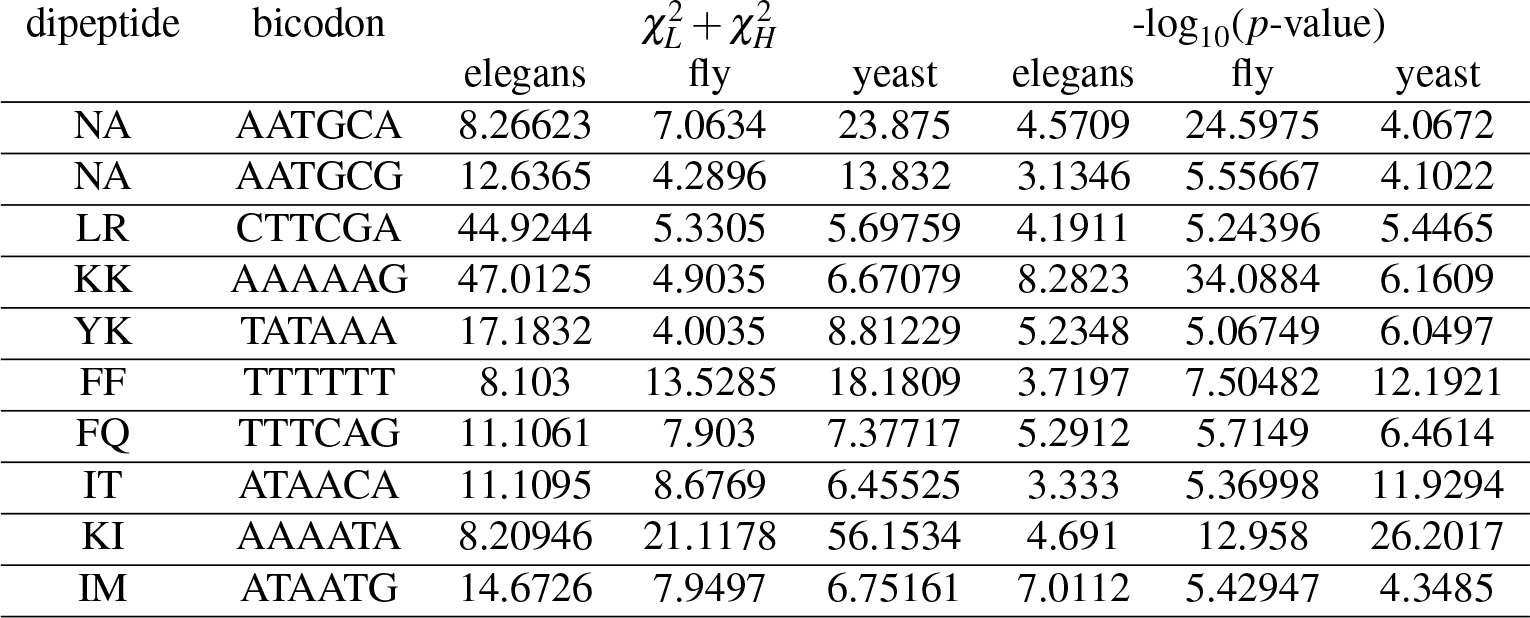
Shared bicodons in *C. elegans*, *D. melanogaster* and *S. cerevisiae* that have high preference for low PA or high PA sequences (*p*-value > 3), but such preference cannot be explained by the codon usage bias (*χ*^2^ >5).

### Supporting evidence for translational attenuation

The statistical analysis above is able to determine which bicodons are associated with low or high abundant proteins. We hypothesized that bicodons associated with low abundant proteins could have a key role in programming translation pauses of the ribosomal machinery. As a proof of principle, we have identified two examples previously reported that can illustrate how bicodons are able to affect translation rates. The first case is the protein sufI in *E. coli*, which does not interact with chaperones, but needs translational pauses for co-translational folding. Zhang *et al.* show that this protein has several transient intermediates^15^. The first multicopper oxidase domain of this protein ends at residue 143. If we consider that the ribosomal exit tunnel can accommodate around 30 residues, the nucleotide sequences responsible for a putative speed attenuation would be downstream, at codons which codify residues 160-180 (Fig. 6A). Fig. 6B depicts the ribosome density profile corresponding to sufI translation, which reveals a high density region around 160-180, i.e., after most of the N-domain has been released from the ribosomal exit tunnel. In addition, around in this position (residue 166) is the highest peak of the degree of folding acquisition measure, Δ*Q*, proposed by Tanaka (Figure 5A of^36^). A naive sequence analysis in this region reveals three Shine-Dalgarno (SD) sequences indicated with asterisks in Fig. 6B. In particular, among nucleotides 486-491 (residues 162-163) there is the hexanucleotide GGTGGA, which has a predicted affinity with the anti-SD sequence of -6.5 kcal mol^−1^ (Fig. 4 of^23^). In addition to this SD sequence, we have found in this region a cluster of bicodons with high pause propensity, i.e., statistically associated with low PA sequences. These bicodons are listed in Fig. 6C. This speed attenuation is not apparent in rate of translation pattern based on the concentration of the tRNA or codon usage, Fig. 6B (bottom panel). Another translational attenuation experimentally tested is one linked to the intermediate 25-28 kDa (around 214-240 residues)^15^. In this case, we have found the same SD sequence that was mentioned above, located 40 residues downstream (more precisely nucleotides 846-851). Again, this hexanucleotide acts together with a small cluster of three bicodons, with high pause propensity, between residues 259 and 276. At this position is also the second highest peak of the folding degree measure (Figure 5A of^36^). These results suggest that bicodons listed in Fig. 6C could be needed for the correct folding of the transient intermediates in *E. coli.*

**Figure 6.**
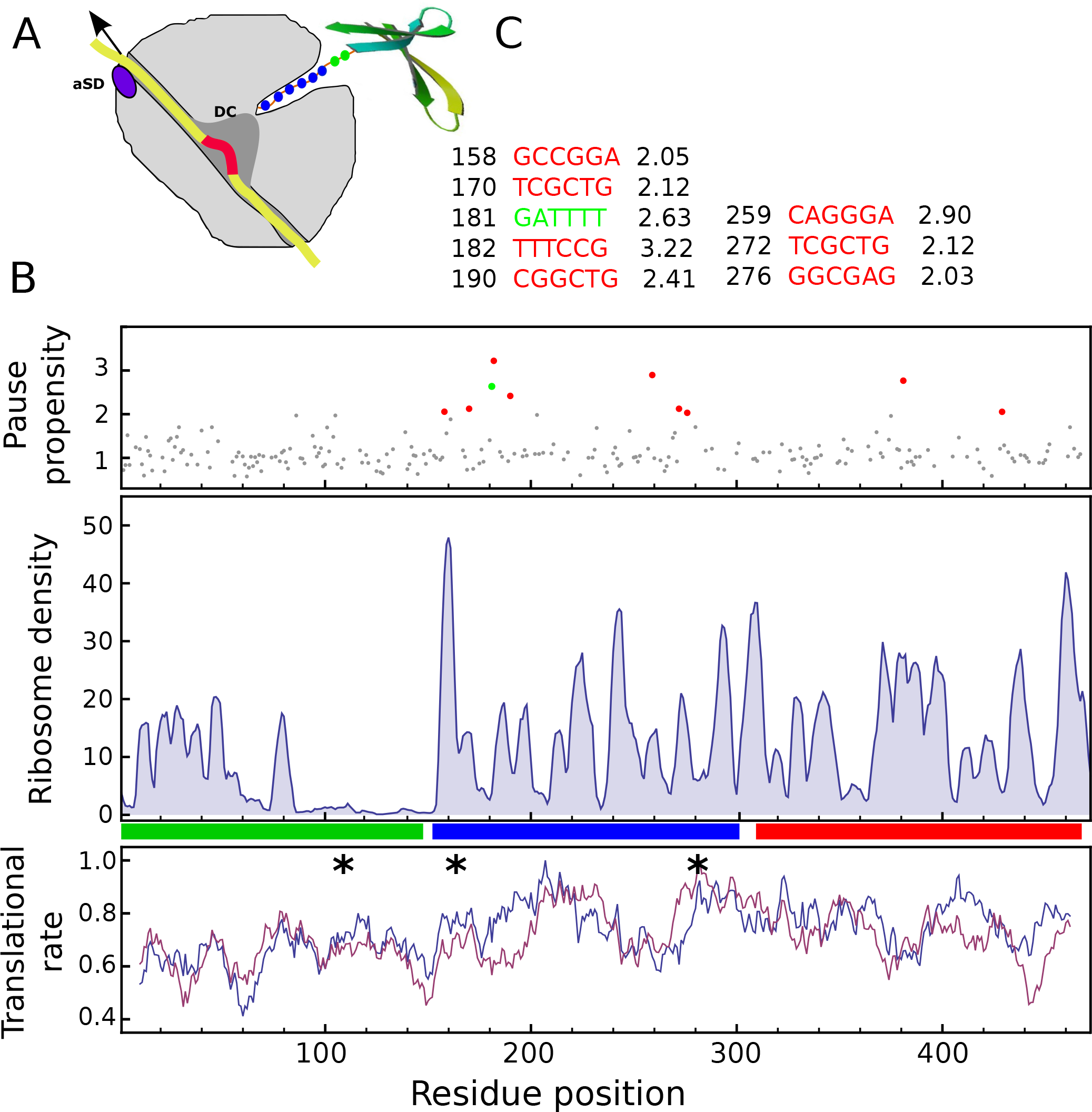
Translational attenuations in the sufIprotein of *E. coli.* (A) Schematic representation of two translational pause mechanisms: anti-SD (aSD) sequence in the 16S RNA can link to SD-like sequences in the transcript. In addition, bicodons with high pause propensity (red) can modulate the translocation rate of tRNAs in the decoding center (DC). (B) Top panel: pause propensity profile of sufI, with two clear cluster of bicodons with high pause propensity (red dots) at 160-180 and at 259-276 residues. These clusters co-localized with SD-like sequences (asterisk). These sequences can be responsible for the peaks in the ribosomal density profile (Middle panel), fact that cannot be explained by the translational rate (Bottom panel) based on codon usage (blue line) or tRNA abundance (red line). (C) Positions, nucleotide sequences and pause propensity of the clusters of bicodons denoted by red and green dots in Fig. 6B.

Further evidence for the role of bicodons in translational pauses is provided by a well studied single polymorphism (SNP) in the gene *MDR1*^37^. This gene encodes the drug-transport pump ABCB1, which transports a variety of drugs from the brain into the blood. Kimchi-Sarfaty *et al.* observed that as a result of a synonymous SNP (rs1045642) the structure and function of the protein are altered with the consequent change in its substrate specificity^16^. The SNP in exon 26 at position 3435 changes the codon ATC to the synonymous ATT, which reduces the codon usage from 47% to 35%. It was argued that the presence of a rare codon affects the timing of co-translational folding and insertion of P-glycoprotein into the membrane. Although it is difficult to consider the ATT codon as rare, it is clear that the SNP alters the timing of the ribosomal transit. We offer here an alternative cause of the translational attenuation; in this sense we have observed that this SNP is also associated with a large change in the propensity pause index *π* of the bicodons. Specifically, bicodon ATCGTG has preference for low PA sequence with *π* = 1.21, while bicodon ATTGTG has preference for high PA sequence *π* = −1.55. This means a change of 178%, almost 8 folds greater than the change in the codon usage. Further, the other synonymous bicodon AT**A**GTG has a even lower pause propensity, *π* = −1.73, in agreement with Kimchi-Sarfaty *et al.* observation that associates to this haplotype a larger decrease in the inhibitory effect^16^. In addition to the SNP above, there are other synonymous SNPs related to human diseases that could be explained by a large change in the pause propensity. Among them we can mention the SNP rs34533956 in the gene CFHR5 which is associated with age-related macular degeneration^38^. In this case, the mutation changes the bicodon GA**C**GTG to GA**T**GTG and associated change in the pause propensity is 183%, while the change in the relative synonymous codon usage is only 13%. Other example corresponds to the SNP rs11615 in the gene ERCC1 which was associated with colorectal cancer^39^, where the pause propensity change is 192%, against a small change in the relative synonymous codon usage (again 13%). These relationship suggest that some pathological synonymous mutations could be understood in terms of the change in the timing needed for co-translational folding programmed by the bicodons.

## Discussion

If we consider an average of three alternative codons for coding each amino acid, there exist more than 1.3 × 10^143^ manners to codify a protein with 300 residues. However, organisms use an insignificant fraction of the number of options offered by the genetic code redundancy. This is due to several constraints operating to optimize many important biological features such as: the expression level^7^, ribosomal proofreading errors^40^, protein solubility^41^, folding accuracy^19,20,42^, protein stability, etc. In this sense, it has been shown that codon usage in *E. coli* is biased to reduce the cost of translational errors^43^. In addition, codons that bind to their cognate tRNA most rapidly are preferentially used in highly expressed genes^44^. It has also been reported a bias in the bicodon usage frequency in several organisms^25^. More recently, Lian *et al.* have identified codons that regulate translation speed in human cell lines^45^. Many other studies agree in the key role of ribosomal pause, coded by codon usage or SD-like sequences, in orchestrating the hierarchical co-translational folding of single domains^15–17,46,47^. In summary, there exist rising evidences that many relevant features, other than the linear sequence of amino acids, are also coded at nucleotide sequence level. These facts should considerably reduce the amount of alternative ways of correctly convey the message from genes to functional proteins, despite the redundancy of the genetic code.

Among the above biological constraints determining the codon usage, we have focused our attention on the translational speed, i.e., the sequential process of protein elongation^4^. Briefly, each proofreading iterative step of this process involves recruitment of the tRNA charged anticodons, tRNA association/dissociation to mRNA, assembly of the residue to the nascent peptide, and translocation of tRNA-mRNA from the A-site to the P-site. Each step has a particular rate and it has been shown that disruption of the interaction between mRNA codon in the A-site from the decoding center is a rate-limiting process^30^. In fact, there are evidences that such rates are codon dependent in *E. coli*^48^. Further contributions to the translational rate, not linked to the tRNAs’ abundance, are the non-Watson-Crick (wobble) interactions. These interactions are usually associated with higher dissociation rates between mRNA and decoding center^49^.

In this paper, we assume the hypothesis that ribosomal pauses are encoded by bicodons, and examine the bicodon frequency usage in nine organisms. We found that many codons have an evident preferential usage in sequences that code for highly abundant proteins, while many others have preference for coding proteins scarcely abundant. The latter bicodons can be understood as short sequences linked to translational pauses. The observed bias cannot been explained by the codon usage in many bicodons. However, the small number of such bicodons found in *E. coli,* where most of bicodon preferences, except for 96 bicodons, can be explained by the codon usage, is worth noting. This clearly contrasts with the other prokaryotes studied here; for example, we have report almost 585 bicodons in *B. subtilis* which are preferentially used to encode either low or high abundant proteins without a codon usage correlate.

The bicodon preference is also found in a plant, a fungus and two invertebrates. Our results indicate that many bicodon preferences are shared by *S. cerevisiae*, *C. elegans* and *D. melanogaster*, and to a lesser extent they are also shared with *A. thaliana* and *B subtilis.* In the case of the mammalian species (*H. sapiens* and *M. musculus*), we found a number of bicodons with high preference comparable with *E. coli*. However, we illustrate with an example of synonymous mutations of clinical relevance, that the exchange of two codons with opposite preferences, even when such preferences are moderate, can alter the translation ribosomal traffic. This example suggests that single mutations that changes the bicodon preference can trigger pathological phenotypes by altering the translational attenuation program of the protein.

Even though the results provided here suggest that some bicodons should regulate translational attenuation, it is important to remark the limitations of the present approach to assign to each bicodon one value of the pause propensity index. The more evident limitation is that protein abundances are not uniquely dictated by a quick translation, transcription levels have also important roles in prokaryotes^7^, while RNAi pathway is a common way to regulate expression in mammalians^50^. These, and other factors, can introduce undesired bias and overshadow some bicodon bias. It is likely that the small number of bicodons with evident preference observed in mammalians is due to the fact that protein abundance is not majorly determined by the bicodon usage, with the consequent poor performance of the method in these organisms. Alternative methods based on the ribosome density profile could overcome these drawbacks. But first it is needed to solve the link between density and ribosomal speed at the nucleotide level, due to the fact that a pause at a given site will stop the transit of many other ribosomes proofreading upstream, increasing artificially the ribosome density of the upstream sequences.

Summarizing, we are here reporting that bicodon usage frequency depends on protein abundance. This preference cannot be explained by the traditional codon usage in many bicodons. This empirical evidence supports the hypothesis that bicodons encode translation pauses. Such a scenario allowed us to contrast this hypothesis in various circumstances where translation rates could be altered. Like the naive codon usage, the bicodon usage can empower novel strategies for rational transcript design that minimize misfolding while simultaneously maximizing co-translational folding for foreign proteins in heterologous hosts.

## Methods

### Data sources

In this work, we have used two kind of data: (i) genome-wide protein abundance across nine model organisms, and (ii) nucleotide sequences associated with proteins indicated above. The absolute protein abundance data from three prokaryotes (*M. aeruginosa B. subtilis* and *E. coli*), one unicellular fungus (*S. cerevisiae*), one plant (*A. thaliana*), two multicellular eukaryotes (*D. melanogaster and C. elegans*) and two mammalians (*M. musculus* and *H. sapiens*) were downloaded from PaxDb web site (http://pax-db.org/) on May 2015^51^. From these comprehensive data sets we have selected two sample of proteins: the 500 most abundant proteins and the 500 less abundant ones, including into our samples only one isoform when more than one are present in the comprehensive data set. As PA distributions are in general biased, i.e., the short proteins should be more abundant than larger ones (see Fig. 7), we have also selected two sets of 500 sequences, but taking into account that the sequence length distribution of both sets were similar (Supplementary Tables S1 and S2). The procedure to sampling sequences with similar length distributions consist in ordering all sequences, corresponding to a given organism, in a PA crescent order. To select the sequences of the low PA samples, we began from the lowest PA extreme of the list of sequences, and we compare 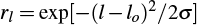 (where *l* is the length of the sequence, *l_o_* and ***σ*** are the mean length and standard deviation, respectively, of the target distribution) with a random number uniformly distributed *r*. If *r_l_* > *r* the sequence is joined to the set of low PA sequences. Then, we test the second sequence in a similar manner and so on, until we have selected 500 sequences. To select the high PA sequence set, we performed the same procedure, but beginning from the end of the sequence list. In Fig. 7 (and also in Supplementary Figs. S3-S10) we have plotted the distributions of whole PA for all organisms used in our study, and the PA and the sequence length distribution of the selected data sets. The nucleotide coding sequences corresponding to the selected proteins were downloaded from Ensembl web sites (four eukaryotes organisms from ftp://ftp.ensembl.org/pub/ and four prokaryotes organisms from http://bacteria.ensembl.org), while *Arabidopsis thaliana* coding sequences were downloaded from www.arabidopsis.org.

**Figure 7.**
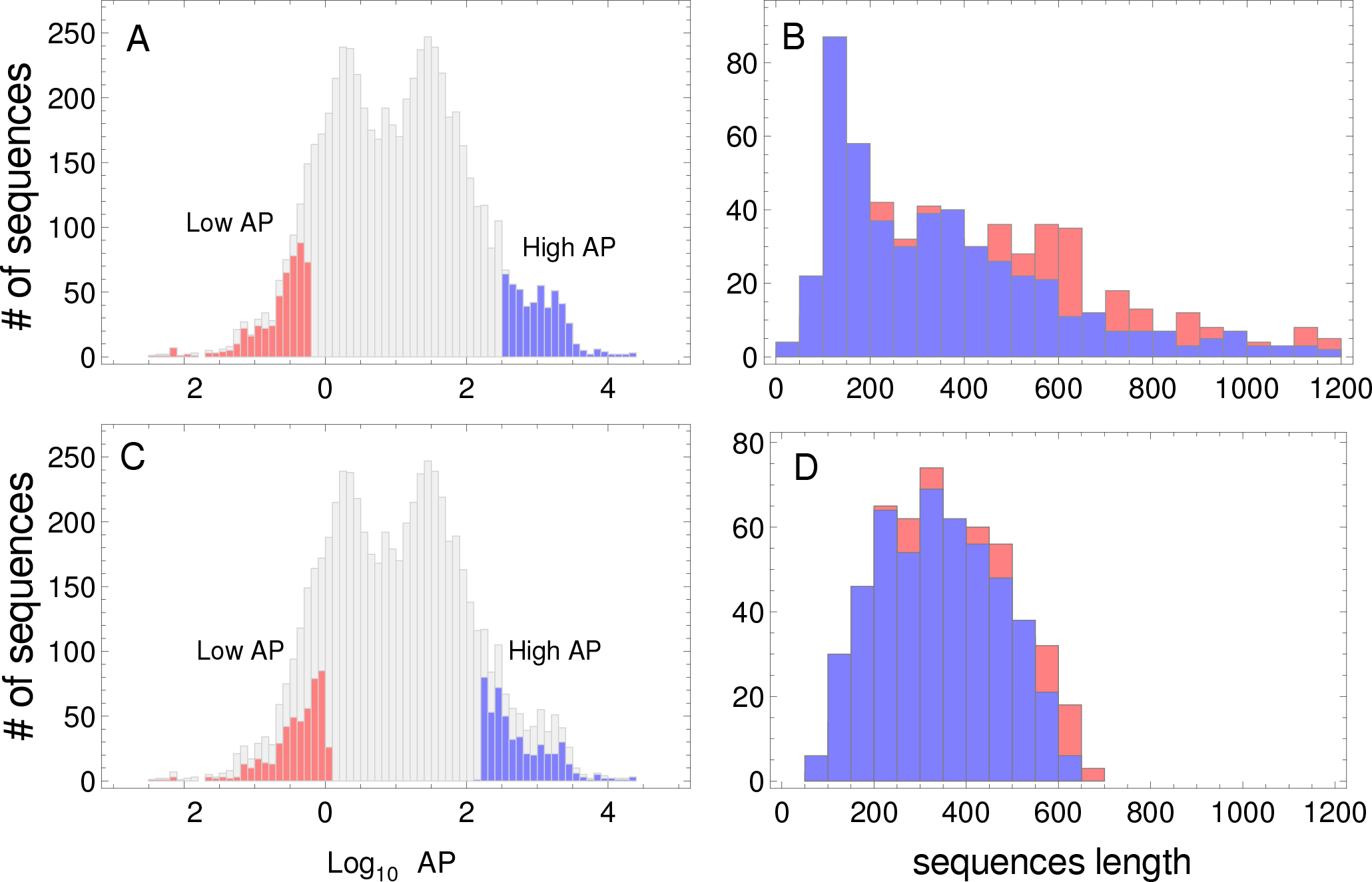
Protein abundance and the sequence length distributions. Protein abundance distributions of the whole dataset of *S. cerevisiae*, lowest and highest PA subsets are indicated in red and blue colors, respectively (A). Sequence length distributions of the subsets of sequences shown in left panel (B). Protein abundance distributions of the whole dataset, the selected low and high PA subsets of sequences used in the study (listed in Supplementary Tables S1 and S2) are indicated in red and blue colors, respectively (C). Sequence length distributions corresponding to the subsets of sequences shown in left panel (D).

We also used ribosome density profiles data of *E. coli* taken from NCBI GEO accession GSE35641^23^. tRNA levels and codon usage of *E. coli* from^44^, and http://www.kazusa.or.jp/codon/, respectively.

### Statistical analyses

The bicodon bias was studied in the context of the low and high PA samples. Basically, we count all consecutive pairs of codons on the same reading frame of the coding sequences belonging to a given sample, which allows us to compute the occurrence of each bicodon *i j* in all sequences of each sample. The index *i* indicates the codon corresponding to P-site, while *j* indicates the one corresponding to the A-site. The occurrence of the codon pair *i j* will be denoted by *oij*. We also compute in the same sample of sequences the number of single codons *f_i_*. Further, we compute expected number of occurrences of each codon pair, as 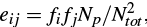, where *N_tot_* is the total number of codons in the set of sequences and *N_p_* is the number of bicodons. Following^34^, we remove the contribution due to the nonrandomness of amino acid pairs by normalizing the former expected values as:

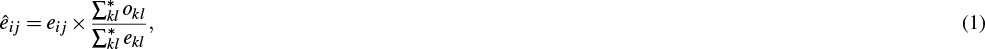

where the * indicates that the sum is only over codon pairs encoding the same amino acid pair encoded for the bicodon *ij*. From the observed and normalized expected bicodon counts recording in a given sample *S*, we compute the residual scores for each codon pair as:

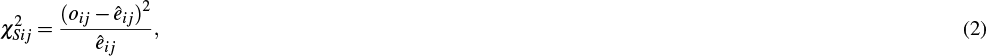

where *S* indicates the sequence samples, i.e., *S* = *L* for low PA sample, or *S* = *H* for high PA sequence sample. Further to use residual scores to test whether the bias in a given codon pair can be explained, or not, by the bias in codons and amino acids, we use the Fisher’s exact test to examine whether a number of occurrences of bicodon 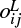, observed in sequences sample associated with low protein abundance are significantly different than the number of occurrences observed in high protein abundance sample 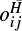. Thus, we construct a 2×2 contingency tables for each bicodon as shown, for an illustrative purpose, in the Supplementary Fig. S11 for the particular case of the bicodon AAGAAG.

Applying the Fisher’s exact test on the right table gives that the observed frequencies of AAGAAG in both samples are significantly different with a p-value of 5.3 × 10^−9352^. In order to compute the binomial coefficients associated with the *p*-value calculation, we approximate the factorial operator with the Stirling’s formula, 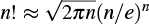for *n* ≥ 25. We performed a similar analysis for all possible bicodons (61 × 64 = 3904), excluding stop:sense bicodons and the stop:stop bicodon. To express the preference degree of a given bicodon for low or high PA sequences we define the pause propensity, *π*, as –*S* log_10_ [*p*-value] where *S* takes value +1, or -1, when the bicodon has preference for sequences with low or with high PA, respectively.

We also use some measures provided by the information theory (IT). The essential IT idea is that of quantify our ignorance associated to a given probability distribution (PD) in a mathematical fashion and formally deal with it. The ignorance associated to a PD {*p*_*i*_} is measured by the Shannon’s entropy^53^ *H*, defined as:

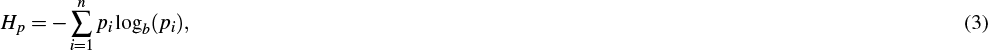

where *b* is the base of the logarithm used; when *b* = 2 the units of entropy are referred to as bits. Hereafter, we work in base 2, and log will be denoted log_2_ in order to simplify the notation. A message (concatenation of symbols of an alphabet) whose symbols have associated a non-uniform distribution will have less entropy than if those symbols had an uniform distribution. This measure is between 0 and 1, and allows us to compare the information content of two PD with different alphabet size. Another IT concept used in this manuscript is the statistical distance between two PDs. In this sense, we use the symmetric version of the Kullback-Leibler divergence measure defined as^54^:

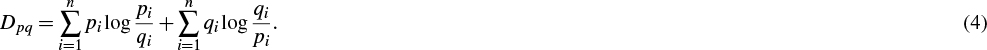

As the alphabet size *n* impacts on the entropy and divergence measures, messages with larger alphabet have associated higher entropy. We found useful to normalize such measures dividing by log(*n*).

## Acknowledgements

We thank Alejandra Carrea and Christina McCarthy for critical reading of the manuscript. LD is member of CONICET (Argentina). This work was partially supported by CONICET, PIP #: 0020.

## Author contributions statement

L.D. conceived and conducted the study, analysed the results and wrote the manuscript.

## Additional information

**Accession codes** GEO accession GSE35641; **Competing financial interests**: The author declare no competing financial interests.

